# Comprehensive preclinical reevaluation of PaBQU and DBQU regimens identifies “lesional liability” in hard-to-treat forms of tuberculosis

**DOI:** 10.64898/2026.09.15.751879

**Authors:** Nicholas D. Walter, Lisa M. Massoudi, Allison A. Bauman, Noalani D. Benedict, Nathan Peroutka-Bigus, Michelle. E. Ramey, Reem Al Mubarak, Samantha Pauly, Firat Kaya, Linda Chaba, Matthew Zimmerman, Debra Flood, David Hermann, Khisimusi Mdluli, David Holtzman, Radojka M. Savic, Jansy Sarathy, Gregory T. Robertson

## Abstract

Murine efficacy models inform advancement of preclinical tuberculosis treatment regimens to human clinical testing. A recent human Phase 2 trial was terminated early because investigational regimens (4-months of PaBQU or DBQU) did not meet the treatment shortening criteria outlined in the Target Regimen Profile (≤ 3 months). We queried whether a factor leading to this early termination may have been “lesional liability,” meaning slow or diminished onset of effect in the caseum of complex lung lesions. We conducted a translational study, comparing the “easy-to-treat” BALB/c mouse, lacking complex lesions, to the “hard-to-treat” C3HeB/FeJ mouse that develops complex human-like lesions. We evaluated traditional and novel pharmacodynamic markers (colony forming units and RS ratio), relapse outcomes and *ex vivo* caseum pharmacokinetics. PaBQU and DBQU were slower to elicit bactericidal activity, RS ratio activity and prevent relapse in the C3HeB/FeJ mouse than the BALB/c mouse. A reference regimen (BPaMZ) had less lesional liability than PaBQU and DBQU. Fewer drugs in PaBQU were projected to achieve target attainment in caseum compared to BPaMZ, particularly early in treatment due to slow accumulation of bedaquiline in caseum. Here, we demonstrated application of a multi-modality pharmacokinetic-pharmacodynamic analysis that compared regimen activity in the hard-to-treat C3HeB/FeJ and easy-to-treat BALB/c mouse models, identifying lesional liability of PaBQU and DBQU. Systematic interrogation of additional diverse regimens is needed to determine the value of preclinical lesional liability as a means of predicting clinical treatment shortening activity.

## INTRODUCTION

Murine efficacy models are employed to evaluate and rank new tuberculosis (TB) treatment regimens prior to testing in humans. It is important to evaluate the concordance of results from murine experimental models with efficacy observed in humans. A recent Phase 2 trial [NCT05971602] tested the efficacy and safety of two novel 4-month regimens that combined one of two nitroimidazole antibiotics (pretomanid [Pa] or delamanid [D]) with a backbone of bedaquiline [B], quabodepistat [Q], and sutezolid [U] (i.e., PaBQU or DBQU). The 4-month trial was designed to inform a second phase in which shorter durations would be tested with the goal of developing a regimen of 3 months or shorter. Because efficacy of the 4-month regimens did not meet pre-defined treatment shortening criteria outlined in the Target Regimen Profile necessary to proceed to evaluation of 3 month and shorter durations, the trial was terminated early. To interrogate the performance of PaBQU and DBQU, we sought to “back-translate” from humans to mice, integrating results from what we consider to be state-of-the-art preclinical tools. This included conventional and novel pharmacodynamic (PD) markers in two preclinical murine relapse models that harbor different *Mycobacterium tuberculosis* (*Mtb*) phenotypes and have differing severity of disease as well as evaluating pharmacokinetic (PK) and PD parameters in caseum.

Contemporary preclinical efficacy evaluation relies primarily on two mouse models that develop markedly different forms of disease: the BALB/c subacute TB infection model and the C3HeB/FeJ chronic infection model.^1,2^ In the BALB/c model, high-dose aerosol deposits ∼10^4^ bacilli in lungs reliably resulting in homogenous non-necrotizing lung lesions with intracellular *Mtb*. The BALB/c subacute model is considered an “easy-to-treat” model both because drug treatment is initiated prior to the onset of adaptive immunity (a point at which *Mtb* is rapidly replicating and highly susceptible to killing) and because the model lacks lesional complexity. In the C3HeB/FeJ model, low-dose aerosol deposits ∼50 to 100 virulent bacilli in lungs, resulting in a highly variable spectrum of disease. A certain proportion of these mice develop overwhelming and rapidly lethal neutrophilic inflammation while others survive with chronic non-necrotic cellular lesions or well-circumscribed caseum-rich necrotic granulomas.^3,4^ Caseating granulomas in the C3HeB/FeJ model recapitulate key features of human TB pathology and harbor an extracellular *Mtb* population.^4–6^ Two factors can make the C3HeB/FeJ mouse a “hard-to-treat” model. First, penetration of certain drugs is slow or limited because necrotic caseum is avascular (requiring drug entry via passive diffusion) and lipid rich (preventing accumulation of lipophobic drugs), and creating temporal pockets of subinhibitory drug concentrations.^7–9^ Second, unlike the rapidly replicating *Mtb* phenotypes found at the start of the BALB/c subacute TB model, *Mtb* populations within C3HeB/FeJ caseating granulomas are slow-growing, metabolically-quiescent and exhibit drug-tolerance.^6,10^ Regimens that under-perform in C3HeB/FeJ mice have been described as having a “lesional liability” attributable to inadequate penetration and accumulation (PK) and/or have limited activity against caseum-adapted bacterial phenotypes (PD). We therefore assessed both caseum PK and PD for PaBQU and DBQU.

In both BALB/c and C3HeB/FeJ mice, efficacy is evaluated based on change in PD markers during treatment as well as relapse outcomes following treatment. Traditionally, PD measures have depended on enumeration of the burden of *Mtb* colony forming units (CFU) in lung tissues. The degree to which drugs or regimens reduce CFU during treatment is described as their bactericidal activity.^11^ A limitation of CFU is that bactericidal activity alone does not consistently predict relapse outcomes (*i.e.,* sterilizing activity).^1,12^ A newer non-culture PD marker is the RS ratio^®^. The RS ratio quantifies ongoing ribosomal RNA (rRNA) synthesis based on the abundance of short-lived precursor rRNA relative to mature structural rRNA.^13^ The RS ratio differs fundamentally from CFU because it measures an aspect of pathogen health rather than pathogen burden. Rather than recapitulate CFU, the RS ratio brings distinct orthogonal information that has been shown to explain variation in relapse outcomes in mice.^13,14^ A limitation of the RS ratio is that protein synthesis inhibitors (including oxazolidinones such as sutezolid included in PaBQU and DBQU) stabilize precursor rRNA,^15–17^ resulting in a rise in the RS ratio unless they are paired with drugs that effectively inhibit ongoing rRNA synthesis.

Since bactericidal activity alone does not consistently distinguish sterilizing activity of different regimens, relapse studies are conducted with both BALB/c and C3HeB/FeJ mice. In the typical relapse design, mice are treated for varying durations then allowed a 3- or more month drug-free holiday before euthanasia ^18–21^ Mice that remain culture positive are classified as having relapsed. Outcomes are typically expressed as an estimation of the time (T) required to prevent relapses in a certain percentage of mice (*i.e.,* 50% [T50], 90% [T90], or 95% [T95]).^22,23^

We compared bactericidal activity, RS ratio activity, and relapse outcomes of PaBQU or DBQU in the easy-to-treat BALB/c model and hard-to-treat C3HeB/FeJ model. We contrasted PaBQU or DBQU with the standard isoniazid [H], rifampin [R], pyrazinamide [Z], and ethambutol [E] regimen (HRZE) and the Simplici-TB regimen,^24^ consisting of B, Pa, Z and moxifloxacin [M] (BPaMZ). Because we observed that PaBQU and DBQU had a slower onset of effect in C3HeB/FeJ mice relative to BALB/c mice, we evaluated caseum PK and PD with caseum-resident *Mtb* phenotypes.

## RESULTS

### Bactericidal activity in BALB/c and C3HeB/FeJ mice

During the first weeks of treatment in BALB/c mice, PaBQU and DBQU displayed greater bactericidal activity than HRZE but lower bactericidal activity than BPaMZ (**Fig 1a-d, Table S1**). Similarly, in C3HeB/FeJ mice, PaBQU and DBQU displayed lower bactericidal activity than BPaMZ (**Fig 1a-d, Table S1**). HRZE was not tested in C3HeB/FeJ mice.

**Figure 1.**
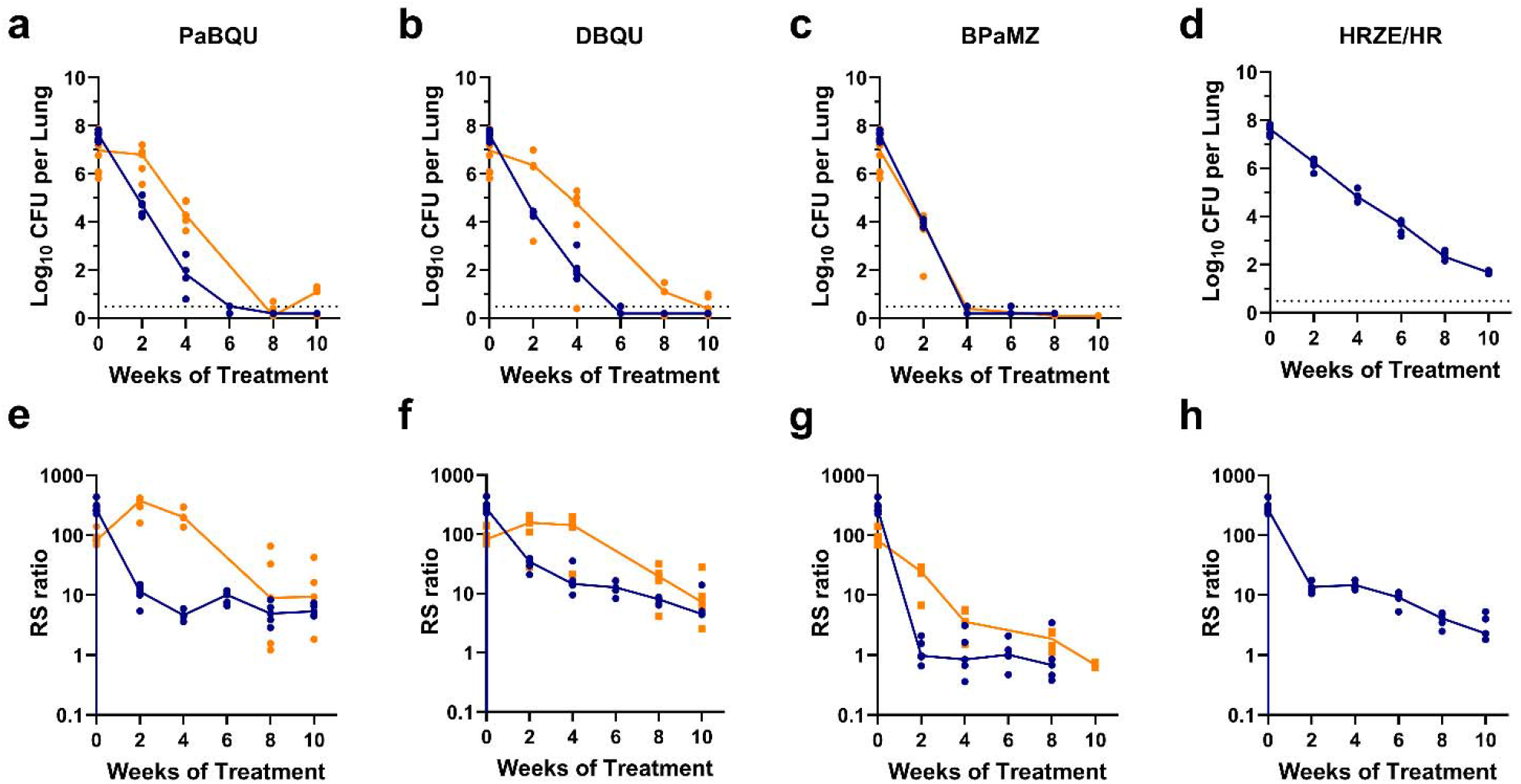
Bactericidal and RS ratio activity of regimens in C3HeB/FeJ mice versus BALB/c mice. *Mtb* Erdman CFU counts (a-d) and RS ratio (e-h) in lungs of C3HeB/FeJ (orange) and BALB/c (blue) mice during the first 10 weeks of treatment with the indicated regimens. Circles represent individual mice and lines connect median values. Mice returning no CFU were plotted at one-half the limit of detection, indicated by the dashed horizontal line.

Comparison between mouse models showed that both PaBQU and DBQU had lower bactericidal activity in the hard-to-treat C3HeB/FeJ model than in the BALB/c model. Specifically, after four weeks of treatment, PaBQU reduced lung CFU by 2.48 log from pre-treatment control in C3HeB/FeJ mice and by 5.80 log in BALB/c mice (*P*=0.0006). After four weeks of treatment, DBQU reduced lung CFU by 2.96 logs in C3HeB/FeJ mice and by 5.46 logs in BALB/c mice (*P*=0.0962). For both PaBQU and DBQU, a longer duration of treatment was required to achieve sterility in C3HeB/FeJ mice than in BALB/c mice (**Fig 1a-b**).

In contrast to PaBQU and DBQU, BPaMZ displayed similar bactericidal activity in C3HeB/FeJ and BALB/c mice (*i.e.,* no significant difference in CFU reduction between the easy-to-treat and hard-to-treat mice). For BPaMZ, the same treatment duration (4 weeks) reduced lung burdens to below lower limits of detection in 4 of 5 mice in both C3HeB/FeJ and BALB/c mice. Although HRZE was not tested in C3HeB/FeJ mice in this study, we have previously observed that HRZE has similar bactericidal activity in C3HeB/FeJ and BALB/c mice.^19^

### RS ratio activity in BALB/c and C3HeB/FeJ mice

In BALB/c mice, PaBQU and DBQU both decreased the RS ratio during the first weeks of treatment, indicating they inhibited rRNA synthesis in the easy-to-treat model (**Fig 1e-h, Table S2**). PaBQU reduced the RS ratio to a significantly greater degree than DBQU (*P*= 0.0004 and 0.0192 at weeks 2 and 4), consistent with previous evidence that pretomanid has greater RS ratio activity.^25^ Both PaBQU and DBQU had lower RS ratio activity than BPaMZ (**Fig 1e-g**).

In C3HeB/FeJ mice, PaBQU and DBQU caused the RS ratio to rise during the first weeks of treatment (**Fig 1e-f**), indicating that the regimens did not effectively inhibit rRNA synthesis in the hard-to-treat model. By contrast, BPaMZ did reduce the RS ratio significantly relative to pre-treatment controls at all time points.

Comparison between mouse models showed differences in RS ratio. At the start of treatment, the median RS ratio was significantly lower in C3HeB/FeJ mice (82.47) than BALB/c mice (270.07) (*P=*<0.0001), consistent with the presence of slowly-replicating and rapidly-replicating *Mtb* phenotypes, respectively. During the first weeks of treatment, PaBQU and DBQU had significantly lower RS ratio activity in C3HeB/FeJ than in BALB/c mice (**Fig 1e-f**). (*P* = 0.0001 and 0.006, respectively). Although the difference between models was not as great as observed for PaBQU or DBQU, BPaMZ also had significantly lower RS ratio activity in C3HeB/FeJ mice than in BALB/c mice (*P=*0.0008 and 0.0223 at weeks 2 and 4, respectively), indicating a modest lesional liability that was not discernable based on CFU alone.

### Relapse outcomes in BALB/c and C3HeB/FeJ mice

PaBQU and DBQU were substantially less effective at preventing relapse in C3HeB/FeJ mice than in BALB/c mice (**Fig 2, Table S3**). Specifically, for PaBQU, the T95 was 138.8 days (95% confidence interval:125.9-151.7 days) in C3HeB/FeJ mice compared with 73.8 days (95% CI: 60.9–86.7 days) in BALB/c mice (*P*<0.0001). For DBQU, the T95 was 157.5 days (95% CI: 144.6–170.4 days) in C3HeB/FeJ mice compared with 83.4 days (95% CI: 70.5–96.3 days) in BALB/c mice (*P*<0.0001). In contrast, BPaMZ, had a statistically significant but more modest difference in T95 between mouse models with a longer T95 in C3HeB/FeJ mice (64.6 days, 95% CI: 51.7–77.5 days) compared with BALB/c mice (44.2 days, 95% CI: 31.3–57.1 days, *P*=0.028).

**Figure 2.**
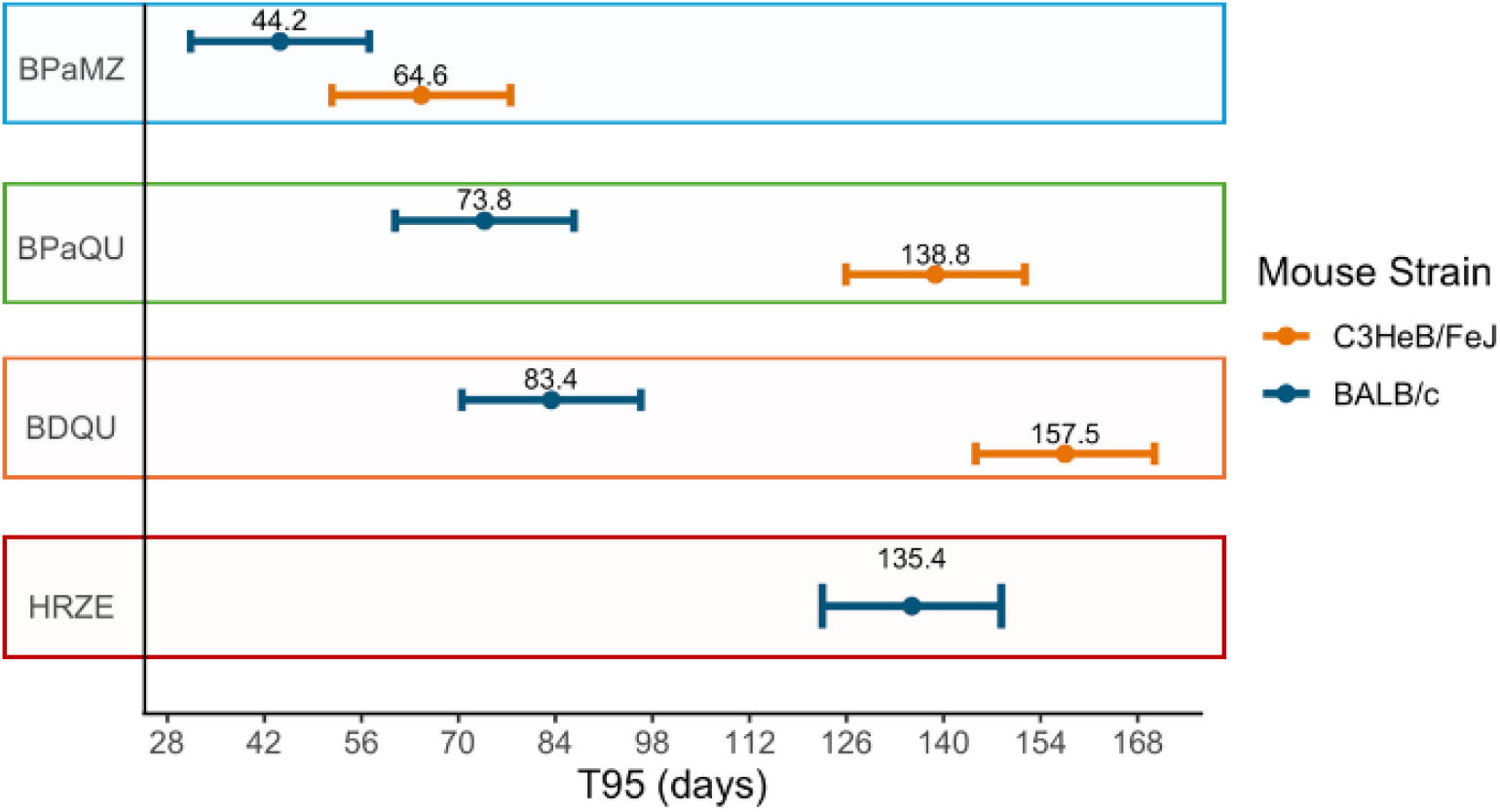
T95 in C3HeB/FeJ versus BALB/c mice. A forest plot of T95 relapse probability in days for C3HeB/FeJ (orange) and BALB/c (blue) mice. The point and whiskers indicate the meedian estimate and 95% confidence interval, respectively.

In BALB/c mice, PaBQU and DBQU were substantially more efficacious than HRZE which had a T95 of 135.4 days (95% CI: 122.5–148.3 days) and less efficacious than BPaMZ which had a T95 of 44.2 days (95% CI: 31.3–57.1 days) (**Fig 2, Table S3**).

### Lesional PK/PD of drugs in the BPaMZ and BPaQU regimens

Because the XBQU regimens under-performed in the hard-to-treat C3HeB/FeJ mouse relative to the easy-to-treat BALB/c mouse, we probed the basis of this lesional liability by evaluating the lesional PK/PD of individual drugs, beginning with bedaquiline.

Consistent with previous evidence that bedaquiline has high RS ratio activity,^26^ testing in the *ex vivo* caseum model^10^ showed that bedaquiline is potent against caseum-resident *Mtb* phenotypes. **Table 1** displays the cMBC_50_ and cMBC_90_, defined as drug concentrations required to kill 50% and 90% of *Mtb* in caseum respectively, and illustrates the potency of bedaquiline relative to other drugs examined in this study. However, consistent with prior evidence that bedaquiline requires weeks to months to reach steady-state concentrations in rabbit lesions,^9^ our testing in the C3HeB/FeJ mice revealed a slow creep of bedaquiline into the central caseous core over repeated doses, such that there is a delay before bedaquiline reaches its cMBC_50_ in inner caseum (**Fig 3a-b**). No drug other than bedaquiline displayed a delay in reaching steady state concentrations in this tissue compartment. Accordingly, we observed similar tissue distribution kinetics for the major circulating bedaquiline metabolite (M2), which is also potent in the *ex vivo* caseum assay and crosses its cMBC_90_ threshold in inner caseum with repeated dosing. We further compared the effect of bedaquiline monotherapy at the accepted human equivalent dose of 25 mg/kg between BALB/c and C3HeB/FeJ mice. Bedaquiline had a rapid and profound effect on CFU and RS ratio in BALB/c mice (**Fig 4a-b**). By day 12, BALB/c mice had 3.15 log reduction in CFU and ∼18-fold decrease in RS ratio. By contrast, consistent with the slow partitioning of bedaquiline into caseous tissue, the effect of bedaquiline on the RS ratio was delayed in C3HeB/FeJ mice. RS ratio declined to a lesser extent in bedaquiline treated C3HeB/FeJ mice with visible necrotic Type I lesions (**Fig 4c**), meaning mice with necrotic lesions were less responsive to bedaquiline and overall had a slower decrease in RS ratio.

**Table 1.** Caseum minimum bactericidal concentration 50 and 90 of drugs included in the PaBQU and BPaMZ regimens. Drug concentrations in µM required to reduce caseum-adapted *Mtb* CFU 50% and 90% as determined via concentration-response testing in the *ex vivo* caseum model.

| Drug | cMBC<sub>50</sub> ($\mu\text{M}$) | cMBC<sub>90</sub> ($\mu\text{M}$) |
| --- | --- | --- |
| moxifloxacin | 0.125 | 1.6 |
| bedaquiline M2 | 1.1 | 4 |
| sutezolid | 1.4 | 16 |
| bedaquiline | 2.1 | 4.7 |
| pretomanid | 3 | 70 |
| sutezolid M1 | 12 | >512 |
| pyrazinamide | 800 | >800 |
| quabodepistat | >512 | >512 |

**Figure 3.**
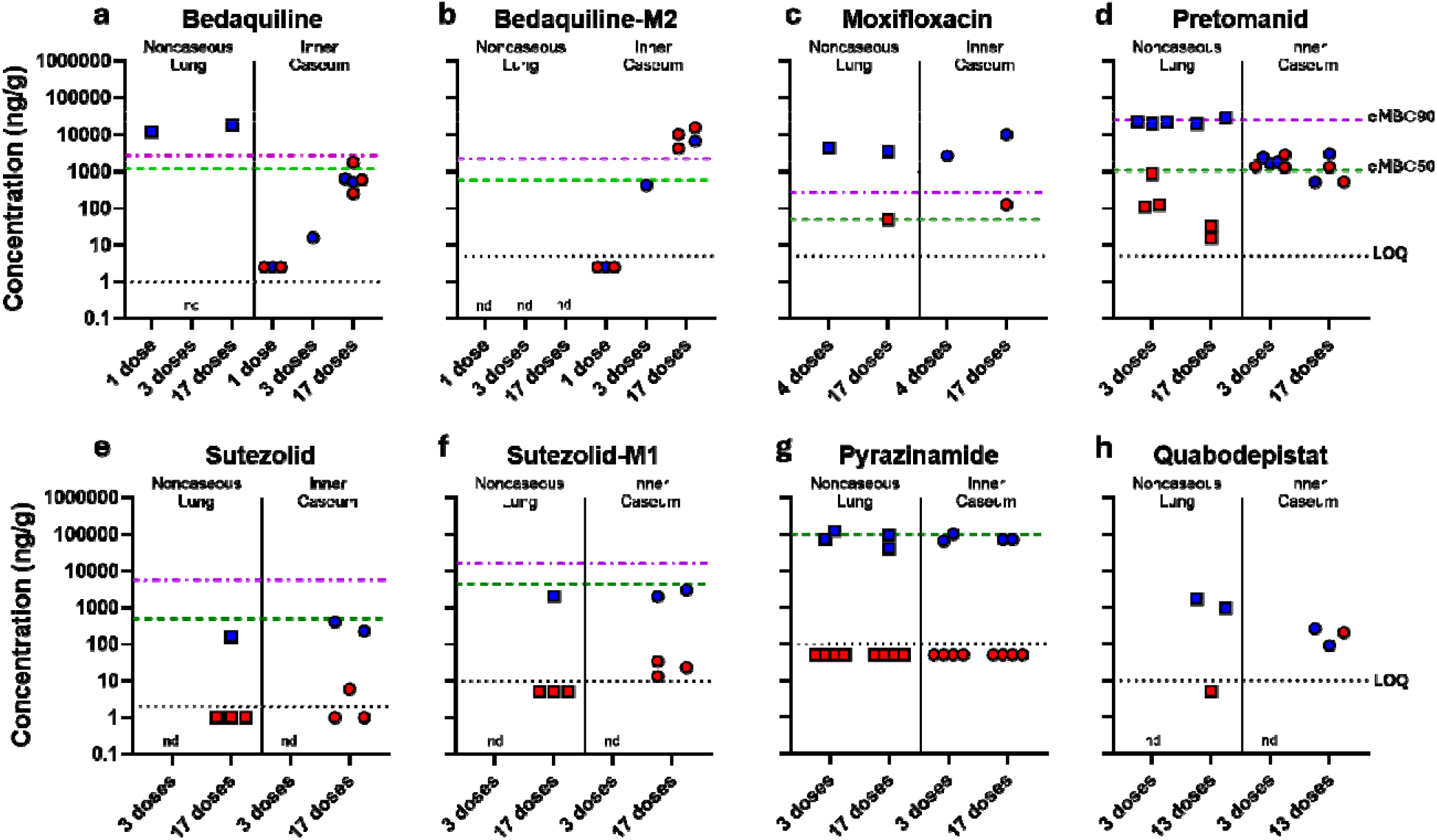
Tissue drug exposure levels in C3HeB/FeJ mice. Mice received the number of indicated doses of each indicated drug, and were necropsied at plasma peak (blue) and trough (red) time points. Drug concentrations were quantified in noncaseous lung compartments (squares) and inner caseum lesion compartments (circles) following precise laser capture microdissection. Drug concentrations are indicated on a log scale. Relevant drug potency measurement cutoffs, cMBC_50_ (green dashed line) and cMBC_90_ (purple dashed/dotted line), which refer to the minimum concentrations required to achieve 50% and 90% bacterial killing in caseum respectively, are indicated with dotted lines. The lower limit of quantification (LOQ) for each drug is similarly indicated (black dotted line). Quabodepistat did not achieve a cMBC_50_ measurement in the concentration range depicted by the y axis.

**Figure 4.**
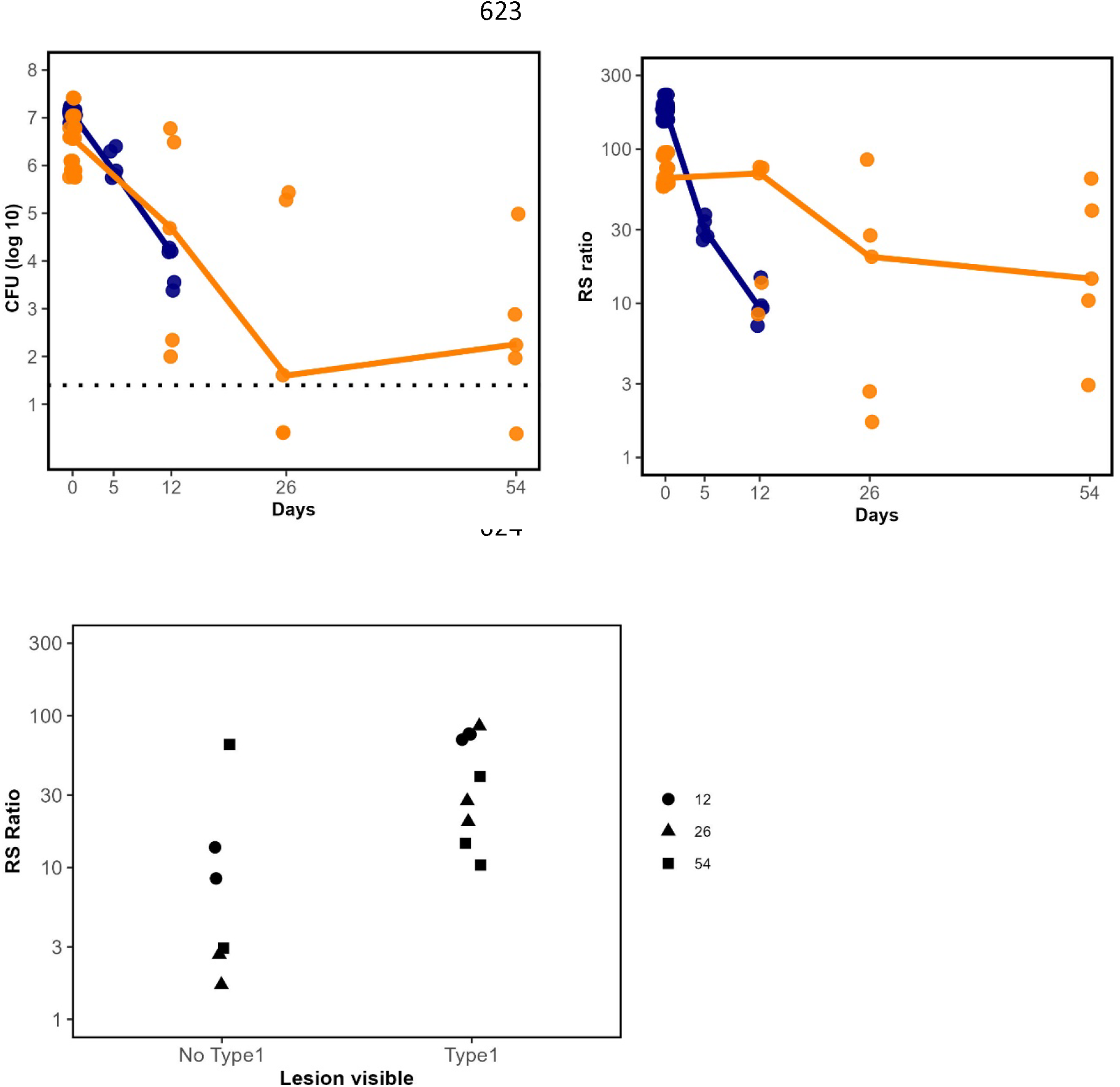
Bactericidal and RS ratio activity of bedaquiline monotherapy in C3HeB/FeJ mice versus BALB/c mice. *Mtb* Erdman CFU counts (a) and RS ratio (b) in lungs of C3HeB/FeJ (orange) and BALB/c (blue) mice during the first 8 weeks of treatment bedaquiline 25mg/kg. Circles represent individual mice and lines connect median values. Mice returning no CFU were plotted at one-half the limit of detection, indicated by the dashed horizontal line. Binning of C3HeB/FeJ mice as having or lacking at least one visible necrotic caseating type 1 lesion at the time of the 12, 26, or day 54 sacrifice relative to their corresponding RS ratio activity.

Testing of other individual drugs in the *ex vivo* caseum model showed a spectrum of caseum potency against nonreplicating *Mtb* bacteria (**Table 1**). The most potent was moxifloxacin which had cMBC_90_ of 1.6 µM. At the opposite extreme, quabodepistat did not achieve either cMBC_50_ or cMBC_90_ at test concentrations up to 512 µM. When tested alone, pyrazinamide is only bactericidal against caseum-resident *Mtb* at high drug concentration (cMBC_50_ of 800 µM). Finally, we compared PaBQU with the closely-related reference regimen, BPaMZ. Since both regimens share the BPa backbone but BPaMZ displayed substantially less lesional liability, we asked what features made moxifloxacin + pyrazinamide better companion drugs than quabodepistat + sutezolid. As highlighted in **Table 1** and **Figure 3**, moxifloxacin reached or exceeded cMBC_50_ concentrations in the caseous compartments of C3HeB/FeJ mice whereas quabodepistat and sutezolid did not. The contribution of pyrazinamide to killing in caseum has been established as combination regimen dependent.^8,19,27^ suggesting it also contributes to controlling caseum-associated *Mtb* phenotypes as a companion drug. These data collectively suggests that companion drugs moxifloxacin and pyrazinamide support the clearance of nonreplicating persistent *Mtb* in necrotic lesions in addition to the efficacy afforded by backbone drugs bedaquiline and pretomanid, driving the superiority of the BPaMZ regimen.

## DISCUSSION

To better understand the PaBQU and DBQU regimens, we evaluated novel and conventional PD markers and relapse outcomes in the easy-to-treat BALB/c and hard-to-treat C3HeB/FeJ mouse models. PaBQU and DBQU displayed considerably lower efficacy in the hard-to-treat C3HeB/FeJ mouse compared with the easy-to-treat BALB/c model, suggesting lesional liability (*i.e.,* diminished efficacy in complex lesions). A closely related reference regimen (BPaMZ) had substantially less lesional liability, demonstrating that achieving comparable efficacy in BALB/c and C3HeB/FeJ models is possible. PK/PD evaluation indicated that a potential explanation for the greater lesional liability of PaBQU relative to BPaMZ is the number of drugs projected to achieve target attainment in caseum, particularly at the start of treatment while bedaquiline is loading. These results suggest the value of comparative efficacy evaluation in easy-to-treat and hard-to-treat mouse models with the use of multiple PD measures and caseum PK/PD analyses.

Human trials show that TB regimen commonly have lower efficacy in patients with advanced pathology than in those with limited disease.^28^ Hard-to-treat patient phenotypes are of particular relevance to the trial of PaBQU and DBQU since 61% of the modified per protocol population reportedly had hard-to-treat disease (*i.e.,* both grade ≥ 2+ sputum smear and radiographically advanced disease).^29^ Comparison of efficacy in the chronic C3HeB/FeJ model versus the subacute BALB/c model provides a basis for estimating efficacy in hard-to-treat phenotypes relative to easy-to-treat phenotypes. Infected BALB/c mice consistently develop uniform diffuse, cellular lesions rather than the complex lesions characteristics of caseum-filled “dirty,” hard-to-treat lesions in human TB.^30,31^ In the easy-to-treat BALB/c model, PaBQU, and DBQU were highly efficacious, reducing time required to cure 95% of mice by 45% and 38% relative to HRZE/HR, respectively. By contrast, infection of C3HeB/FeJ mice results in a variable spectrum of lung pathology. Some mice develop uncomplicated cellular lesions similar to those observed in the BALB/c subacute model. Others develop well-circumscribed, avascular, necrotic lesions with a lipid-rich, hypoxic caseus core,^4,32^ that have important similarities with advanced human pathology. A central finding of this project is that PaBQU, and DBQU had substantially lower efficacy in C3HeB/FeJ mice than in BALB/c mice. Specifically, the T95 of PaBQU and DBQU was nearly twice as long in C3HeB/FeJ mice compared with BALB/c mice. This under-performance in models with advanced pathology has been described as lesional liability.

The reference regimen used in these studies (BPaMZ) demonstrated that not all regimens have a high degree of lesional liability. We found that BPaMZ was highly efficacious in both the hard-to-treat and easy-to-treat mouse models. The T95 of BPaMZ was only marginally longer in C3HeB/FeJ mice with confidence intervals overlapping the T95 in BALB/c mice. In C3HeB/FeJ mice, BPaMZ was more efficacious than PaBQU, and DBQU, curing 95% of mice in less than half the treatment duration.

Our observation that BPaMZ and PaBQU have substantially different efficacy in C3HeB/FeJ mice provided an opportunity to probe the basis of lesional liability by evaluating the lesional PK/PD of individual drugs. The drugs shared by BPaMZ and PaBQU (bedaquline and pretomanid) both displayed potent activity against metabolically quiescent, non-replicating caseum *Mtb* phenotypes. Both bedaquiline and pretomanid ultimately accumulate at concentrations predicted to achieve target attainment in caseum. However, bedaquiline is “slow to load,” taking weeks to achieve effective concentrations in caseum.^9^ Two of the drugs included in these regimens (quabodepistat and pyrazinamide) are unlikely to contribute appreciably to lesional activity since they lack potency against caseum *Mtb* phenotypes and do not reach target attainment concentrations in caseum. PK/PD evaluation showed moxifloxacin was more potent than sutezolid against caseum phenotypes and reached target attainment concentrations. Although sutezolid had activity against caseum *Mtb* phenotypes when tested in the *ex vivo* caseum model, neither sutezolid nor its M1 metabolite are predicted to reach target attainment concentrations in caseum *in vivo.* Since bedaquiline is slow to reach effective concentrations in caseum, BPaMZ includes two drugs (pretomanid and moxifloxacin) that are effective in caseum at the start of treatment, with three drugs being effective once bedaquiline achieves steady-state levels in caseum later in treatment. By contrast, PaBQU includes one drug (pretomanid) that is moderately effective in caseum at the start of treatment, increasing to two drugs providing lesion coverage later in treatment once bedaquiline achieves steady-state levels in caseum.

As PD markers, we used both the traditional culture-based measure of bacterial burden (CFU) and a novel measure of pathogen health (RS ratio). Both CFU and RS ratio revealed the lesional liability of DBQU and PaBQU in different ways. CFU showed that the bactericidal activity of DBQU and PaBQU was lower in C3HeB/FeJ mice than in BALB/c mice, whereas the bactericidal activity of BPaMZ was similar in C3HeB/FeJ and BALB/c mice. As highlighted in the PK/PD analyses above, the observation that BPaMZ lacked bactericidal lesional liability is potentially attributable to the presence of a second caseum-active drug (moxifloxacin) at the start of treatment while bedaquiline is loading into caseum. RS ratio results also highlighted lesional liability for PaBQU and DBQU. Concordant with historical observations,^15–17^ oxazolidinones are known to increase the RS ratio^26^ unless other drugs in the regimen effectively inhibit ongoing rRNA synthesis. In the BALB/c mouse, PaBQU and DBQU rapidly decreased the RS ratio, suggesting that tested oxazolidinones quickly penetrated macrophages and inhibited rRNA synthesis. PaBQU had notably greater RS ratio activity than DBQU which is consistent with the known greater RS ratio activity of pretomanid relative to delamanid and consistent with the trend towards a shorter T95 with PaBQU. By contrast, in the C3HeB/FeJ mouse, the RS ratio rose significantly after 2 and 4 weeks of treatment before declining at week 8. This effect is consistent with the slow arrival of bedaquline (a drug with strong RS ratio activity^13,26^) in lesions. Finally, for BPaMZ, the RS ratio decreased more slowly in C3HeB/FeJ mice than in BALB/c mice, revealing a limited degree of lesional liability that was not apparent based on CFU. This is consistent with the PK/PD analysis and with the trend towards longer T95 in C3HeB/FeJ mice. The finding that lesional liability was discernable based on CFU and RS ratio results early in treatment is important because these pharmacodynamic results can be obtained much more quickly and at lower cost with fewer mice than conventional studies used to estimate T95.

These results support the use of a stepwise regimen evaluation pathway with a focus on lesional liability. This begins with *ex vivo* quantification of the potency of individual drugs against caseum-resident *Mtb* phenotypes (*i.e.,* cMBC50 and cMBC90). This lesional potency information is combined with lesional PK to estimate steady-state lesion coverage for individual drugs and full combinations while also considering the time drugs require to reach steady state concentration in caseating lesions. Lesional liability is then assessed empirically *in vivo* via head-to-head comparison of the easy-to-treat BALB/c and hard-to-treat C3HeB/FeJ mouse models based on PD markers (CFU and RS ratio) early in treatment and eventual relapse results (quantified as T95). To validate the relevance of the C3HeB/FeJ mice relative to BALB/c mice comparison to efficacy in human disease, it will be necessary to systematically interrogate a number of diverse regimens for which clinical efficacy is established. In the current proof-of-concept case, preclinical lesional liability seems concordant with clinically observed bactericidal activity. Specifically, in SimpliciTB, 8-week culture conversion was achieved in 122 (84%) of 145 participants treated with four months of BPaMZ versus 70 (47%) of 148 participants treated with four months of HRZE. In the XBQU trial, 8-week culture conversion was achieved in 6 (38%) of 16 participants treated with PaBQU and 6 (40%) of 15 participants treated with HRZE.

This study has several limitations. This was a secondary analysis, leveraging results from separately designed relapse studies. Consequently, there was a difference in which DBQU and PaBQU were dosed 7 days a week in the BALB/c study, but only 5/7 days per week in the C3HeB/FeJ study. As a sensitivity analysis, we conducted a 1-month comparison of 5/7 versus 7/7 dosing with PaBQU and DBQU. As summarized in Supplementary Information Section 1, the dosing schedule did not affect bactericidal activity. For PaBQU, 7/7 dosing yielded greater RS ratio activity than 5/7 dosing in the BALB/c mouse but this effect was diminutive relative to the RS ratio differences observed between C3HeB/FeJ and BALB/c mice. We conclude that this does not affect our central findings regarding lesional liability. The role of lesional liability will be more convincingly demonstrated by forward-translation in which these preclinical tools predict the clinical efficacy of regimens currently in clinical trials, work that is ongoing through the Preclinical Design and Clinical Translation of TB Regimens (PReDiCTR) Consortium (https://www.predictrtb.org/consortium).

Synthesis of start-of-the-art preclinical evaluation tools articulated the concept of lesional liability as a measure that has potential to inform regimen progression. Our results suggest that lesional liability is identifiable during preclinical testing via the integration of a suite of tools. We propose that stepwise evaluation of candidate regimens harnessing these state-of-the-art tools may improve prediction of clinical results in the future.

## MATERIALS & METHODS

### Bacterial strains

*Mtb* strain Erdman (TMCC 107) was used *in vitro* to infect mice. *Mtb* strain HN878 was used to generate rabbit caseum.^10^

### Murine Infection Models

All procedures and protocols for infecting mice with *Mtb* and subsequent drug treatments were approved by the Colorado State University Institutional Animal Care and Use Committee (IACUC) (Approved protocol number: KP 5172). Mice were housed in a certified animal bio-safety level III (ABSL-3) facility. Water and mouse chow were provided *ad libitum*. Specific pathogen-free status was verified through sentinel animals and/or by environmental monitoring of facility ABSL3 exhaust filters for opportunistic pathogens. Mice were block randomized after aerosol infection and distributed into the different treatment arms. Infected mice were observed daily and weighed up to two times per week, due to the increased incidence of morbidity and mortality associated with clinical TB disease in the subacute BALB/c and the chronic C3HeB/FeJ mouse models. Any mice exhibiting clinical symptoms of illness were humanely euthanized.

### Drugs and experimental compounds

Quabodepistat and spray-dried delamanid were provided by Otsuka. All other drugs were from commercial sources.

### Drug formulation

Delamanid (D) and quabodepistat (Q) were formulated in 5% gum Arabic. Sutezolid (U) was formulated in 10% polysorbate 80, 40% polyethylene glycol 400 solution. Bedaquiline fumarate (B) was formulated in an acidified 20% 2-hydroxypropyl-β-cyclodextrin solution. Pretomanid (Pa) was formulated in a 10% 2-hydroxypropyl-β-cyclodextrin solution. Moxifloxacin hydrochloride and pyrazinamide were formulated in sterile water. Isoniazid (H), rifampin (R), pyrazinamide (Z), ethambutol (E) were prepared as described (PMID: 39345140).

### BALB/c subacute TB high dose aerosol model

Female 6- to 8-week-old BALB/c mice were from Jackson Laboratories (Bar Harbor, ME). Mice were aerosol-infected using an inhalation exposure system (Glas-Col, Terre Haute, IN) with Mtb Erdman sub-cultured in Middlebrook 7H9 broth supplemented with 0.2% glycerol, 10% acid-albumin-dextrose-catalase (ADC) and 0.05% Tween 80 and adjusted to an optical density at 600 nm (OD600) of 0.8. This resulted in the deposition of 4.56 log10 CFU in lungs of each mouse one-day after aerosol. Nine mice were humanely killed 11 days post aerosol (D0) to determine CFU lung burdens at the start of treatment. The remaining mice received one of four different treatments with combinations of drugs at the following doses (in mg/kg body weight): bedaquiline (25), pretomanid (50 [Pa_50_] or 100 [Pa_100_], as indicated below), spray-dried delamanid, delamanid (6), quabodepistat, Q (9), moxifloxacin hydrochloride, M (100), sutezolid, U (50), isoniazid, H (10), rifampin, R (10), pyrazinamide, Z (150), ethambutol, E (100). R was administered by gavage in 0.2 mL volume, followed >2 h later by companion drugs or drug combinations (e.g., H or HZE). For combination studies HZE was combined immediately before administration and given as a single oral gavage in 0.2 mL. HRZE was given 5 of 7 days per week with mice receiving only HR after the first 2 months of intensive therapy with all four drugs (i.e., HRZE). BPa_100_MZ was included as a reference benchmark control (PMID: 34006838) and was administered 5 of 7 days per week. The two experimental study arms included Pa_50_BQU, and DBQU. Each drug for the experimental arms was given individually in 0.2 mL volume by gavage 7 days per week. Daily administered doses were separated by ≥ 1 h.

### C3HeB/FeJ chronic TB low dose aerosol model

Female 8- to 10-week-old C3HeB/FeJ mice from Jackson Laboratories (Bar Harbor, ME) were aerosol-infected using an inhalation exposure system (Glas-Col, Terre Haute, IN) with a previously calibrated *Mtb* Erdman frozen aliquot that was thawed and diluted to an average titer of ∼1×10^6^ CFU/mL in sterile water prior to infection. The average bacterial load in lungs of each mouse one-day after low-dose aerosol was 1.94 log10 CFU. Eight mice were humanely killed 56 days post aerosol (D0) to determine CFU lung burdens at the start of treatment. Mice received one of three experimental treatment arms (BPa_100_MZ, Pa_50_BQU, or DBQU) at the doses listed above. D or Pa was administered by gavage in 0.2 mL volume, followed 4 h later by BQU combined immediately before administration and given as a single oral gavage in 0.2 mL. BPa was combined immediately before administration and given as a single oral gavage in 0.2 mL followed 4 h later by MZ, which was combined immediately before administration and given as a single oral gavage in 0.2 mL.

A separate bedaquiline monotherapy study was conducted in 8-10-week-old female C3HeB/FeJ mice following the above infection protocol. The average bacterial load in lungs of each mouse one-day after low-dose aerosol was 1.70 log10 CFU. Bedaquiline was administered for 2 months, 5 of 7 days per week in 0.2 mL volume by oral gavage at 25 mg/kg.

### Drug efficacy experiments and relapse

Efficacy determinations were based on lung CFU counts from whole lungs (C3HeB/FeJ mice) or 2/3rds (BALB/c mice) of the lung (by weight). Tissues were aseptically harvested and flash frozen in 7mL Bertin Precellys tubes (CKmix50_7mL P000939-LYSK0A) under liquid nitrogen. Samples were thawed, homogenized (Precellys, Bertin Instruments, Rockville, MD) and serially diluted in PBS with 10% [w:v] bovine serum albumin. Portions of the homogenates from animals on treatment were plated for CFU on 7H11-OADC agar (i.e., Middlebrook 7H11 agar plates supplemented 0.2% [v:v] glycerol, 10% [v:v] oleic acid-albumin-dextrose-catalase (OADC) supplement, and 0.01 mg/mL cycloheximide, and 0.05 mg/mL carbenicillin) further supplemented with 0.4% [w:v] activated charcoal (7H11 charcoal agar) to help counteract drug carryover artifacts (PMID: 37791784, PMID: 26503656, PMID: 39345140). Lung homogenates from the relapse arms (i.e., following a 12 week [3 month] drug holiday) were plated on 7H11-OADC agar without charcoal, but further supplemented with 25 mg/L polymyxin bedaquilinine and 20 mg/L trimethoprim to help prevent potential sample loss due to contamination. Colonies were enumerated after at least 28 days of incubation at 37°C and plates were incubated for ≥ 6 weeks to ensure all viable colonies were detected. Mice were euthanized by CO_2_ inhalation followed by cervical dislocation, a method approved by the IACUC at Colorado State University.

### Methods to estimate of time to cure 95% of mice

The time required to cure 95% of mice (T95) was derived analytically using logistic regression models.^33^ Separate models were fitted to data from two relapsing mouse model (RMM) studies conducted in BALB/c and C3HeB/FeJ mice, which evaluated a common set of treatment regimens and were therefore well suited for direct strain comparison. In each model, binary relapse status (relapsed vs. not relapsed) was modeled as the outcome, with treatment duration (days) and regimen included as covariates. T95 was calculated as the treatment duration corresponding to a predicted relapse probability of 0.05. Ninety-five percent confidence intervals (95% CIs) for T95 were estimated using the delta method.^23,33^

#### *Ex vivo* caseum bactericidal activity assay

The bactericidal activity of TB drugs against nonreplicating *M. tuberculosis* in ex vivo rabbit caseum was conducted as described previously.^10,34^ Briefly, caseum excised from the cavities of TB-infected New Zealand White rabbits was 3-fold diluted, homogenized and added to 96-well plates spotted with the appropriate concentration of each test agent. Each drug was evaluated in the concentration range of 0.5 – 512 µM, with the exception of pyrazinamide, which was evaluated up to the test concentration 2,048 µM in order to reflect the high pyrazinamide clinical dose and corresponding high systemic exposure. After 7 days of drug exposure, each well was sampled and serially diluted prior to plating on supplemented 7H11 agar containing 0.4% activated charcoal. Colony-forming units (CFU) were enumerated 4 weeks later. A DMSO-only control well was included in each assay.

#### RNA extraction

Owing to heterogeneity in pathology/disease burden, RNA was extracted from the C3HeB/FeJ lung homogenate immediately after homogenization at a ratio of 2 to 1 CAMM-RPS buffer (GTC-TCEP) to homogenate. RNA was extracted from flash frozen superior and middle flash frozen lung lobes from BALB/s mice in CAMM-RPS, as described previously (PMID: 37905920).

#### RS ratio

Details on the RS ratio pharmacodynamic measure were described previously (PMID: 34006838).

### Plasma LC-MS/MS

Bedaquiline, pretomanid, delamanid, quabodepistat, moxifloxacin, sutezolid, isoniazid, rifampicin, pyrazinamide and ethambutol were purchased from Sigma-Aldrich. Deuterated internal standards for each analyte were obtained from Toronto Research Chemicals. Drug free K2EDTA plasma and lungs were obtained in-house and from BioIVT for use as blank matrices to build standard curves. Neat 1 mg/mL DMSO stock for analytes were serial diluted in 50/50 acetonitrile water to create standard curves and quality control spiking solutions. Spiked matrix standards and QCs were created by adding 10 µL of spiking solutions to 90 µL of drug free plasma. Extraction was performed for standards, QCs, and study samples by adding 200 µl of 1:1 acetonitrile (ACN)/methanol (MeOH) containing deuterated internal standards to 20 µl of plasma. Samples were vortexed for 5 minutes at room temperature and 100 µl of supernatant were transferred to a 96 well plate for injection. Ten µl of 75 mg/ml ascorbic acid was added to all wells to stabilize rifampicin. For isoniazid, 1:1 2% trans-cinnamaldehyde in MeOH was added, and samples were vortexed for 30 minutes to derivatize isoniazid.

LC/MS-MS analysis was performed on a Sciex Qtrap 6500+ triple-quadrupole mass spectrometer coupled to a Shimadzu Nexera X2 UHPLC system to quantify each drug in plasma. Chromatography was performed on a Phenomenex Luna Polar C18 (2.1×100 mm; particle size, 3 µm) using a reverse phase gradient elution with aqueous. Milli-Q deionized water with 0.1% formic acid (FA) was used for the aqueous mobile phase and 0.1% FA in ACN for the organic mobile phase. Normal phase was used for quantifying ethambutol and performed on a Waters Xbridge Premier BEH amide (2.1×150mm; particle size, 2.5 µm) column. Multiple-reaction monitoring (MRM) of precursor/fragment transitions in electrospray positive-ionization mode was used to quantify the analytes. MRM transition of 555/58, 360/175, 535/352, 457/176, 402/358, 354/312, 252/80, 823/791, 205/116 and 124/81, were used for bedaquiline, pretomanid, delamanid, quabodepistat, moxifloxacin, sutezolid, isoniazid, rifampicin, pyrazinamide and ethambutol respectively. Sample analysis was accepted if the concentrations of the quality control samples were within 20% of the nominal concentration. Data processing was performed using Analyst software (version 1.7.2; Sciex).

### Laser capture microdissection

Twenty-five µm thick tissue sections were cut from mouse lung using a Leica CM 1860UV (Buffalo Grove, IL) and thaw-mounted onto 1.4 µm thick Leica PET-Membrane Frame Slides (Buffalo Grove, IL) for laser capture microdissection. Tissue sections were immediately stored in sealed containers at -80°C. Adjacent 10 µm thick tissue sections were thaw-mounted onto standard glass microscopy slides for H&E. Cellular, necrotic (caseum), uninvolved lung lesion areas for lung totaling 3 million µm^2 were dissected from between 1 to 2 serial biopsy tissue sections using a Leica LMD6 system (Buffalo Grove, IL). The total tissue volume of each pooled sample was determined based on the surface area of the pooled sections and the 25 µm tissue thickness. Areas of interest were identified optically from the brightfield image scan and by comparison to the adjacent H&E reference tissue. Pooled dissected tissues were collected into 0.25 mL standard PCR tubes and immediately transferred to the -80°C.

Neat 1 mg/mL DMSO stocks for all compounds were diluted serially in 50/50 ACN:H2O to create standard curves and quality control spiking solutions. 2 µL of neat spiking solutions were added to 2 µL of tissue homogenate were combined prior to extraction. 2 µL of ACN:H2O and 2 µL of PBS were added to the dissected study samples. Extraction was performed by adding 50 µL of extraction solution ACN/MeOH (1/1). Extracts were vortexed for 5 minutes and centrifuged at 10,000 rpm for 5 minutes. 50 µL of supernatant was transferred for LC/MS-MS analysis. See HPLC-mass spectrometry section for LC/MS-MS parameters.

## Acknowledgements

This work was supported by the Gates Foundation under grant ID number INV-009105 to GR ‘TB Drug Accelerator: TB mouse in vivo models.’ The conclusions and opinions expressed in this work are those of the authors alone and shall not be attributed to the Foundation. We acknowledge Laura Via and Clifton Barry 3rd at the National Institute of Health for rabbit caseum, Nicole Frahm for comments on the manuscript, and the staff of the Laboratory Animal Resources at Colorado State University for animal care.

