## Supplemental information for "Comprehensive preclinical reevaluation of PaBQU and DBQU regimens identifies “lesional liability” in hard-to-treat forms of tuberculosis"

### **TABLE S1**. Lung CFU counts in BALB/c or C3HeB/FeJ mice.

CFU were assessed before and after treatment for the indicated number of weeks of therapy with the listed regimens.

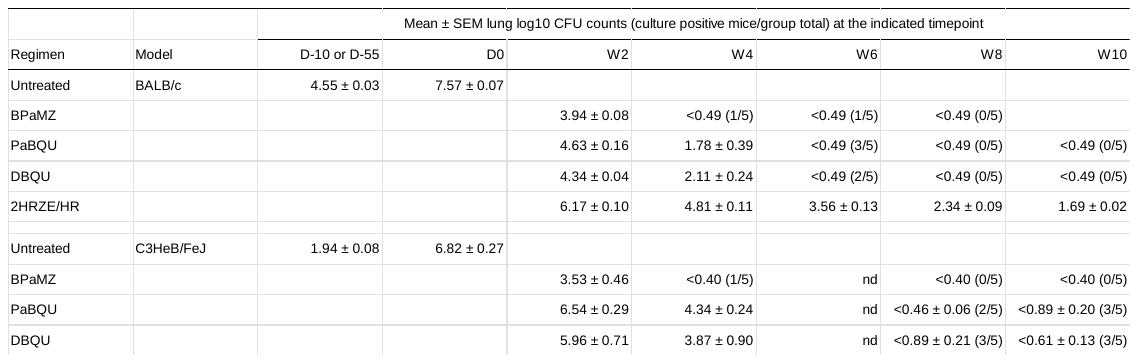

D-10, day -10; W2, week 2, etc..; nd; not determined

### **TABLE S2**. Lung RS ratio in BALB/c or C3HeB/FeJ mice.

RS ratio was assessed before and after treatment for the indicated number of weeks of therapy with the listed regimens.

| Mean±SEM lung RS ratio at the indicated timepoint | | | | | | | |
| --- | --- | --- | --- | --- | --- | --- | --- |
| Regimen | Model | D0 | W2 | W4 | W6 | W8 | W10 |
| Untreated | BALB/c | 305.6 ± 26.2 |  |  |  |  |  |
| BPaMZ |  |  | 1.2 ± 0.3 | 1.3 ± 0.5 | 1.1 ± 0.3 | 1.2 ± 0.6 | nd |
| PaBQU |  |  | 11.0 ± 1.6 | 4.6 ± 0.4 | 9.3 ± 1.0 | 5.3 ± 1.0 | 5.7 ± 0.6 |
| DBQU |  |  | 31.7 ± 3.1 | 17.9 ± 4.5 | 13.0 ± 1.6 | 7.6 ± 0.5 | 6.4 ± 1.9 |
| 2HRZE/HR |  |  | 14.4 ± 1.4 | 15.2 ± 1.1 | 8.8 ± 1.0 | 4.0 ± 0.5 | 3.1 ± 0.7 |
| Untreated | C3HeB/FeJ | 89.5 ± 7.6 |  |  |  |  |  |
| BPaMZ |  |  | 22.5 ± 4.1 | 3.9 ± 0.8 | nd | 1.8 ± 0.3 | 0.7 ± 0.1 |
| PaBQU |  |  | 325.4 ± 45.8 | 224 ± 30.7 | nd | 22.2 ± 12.4 | 15.4 ± 7.2 |
| DBQU |  |  | 135.9 ± 31.3 | 135.6 ± 31.3 | nd | 18.6 ± 4.4 | 10.2 ± 4.4 |

D-10, day -10; W2, week 2, etc..; nd; not determined

### **TABLE S3**. Relapse proportions in BALB/c and C3HeB/FeJ mice.

The fraction of mice with microbiologic relapse 12-weeks after completion of various durations of BPaMZ, PaBQU, DBQU and 2HRZE/HR is shown for BALB/c and C3HeB/FeJ mice. Results for 2HRZE/HR were not available in the C3HeB/FeJ model.

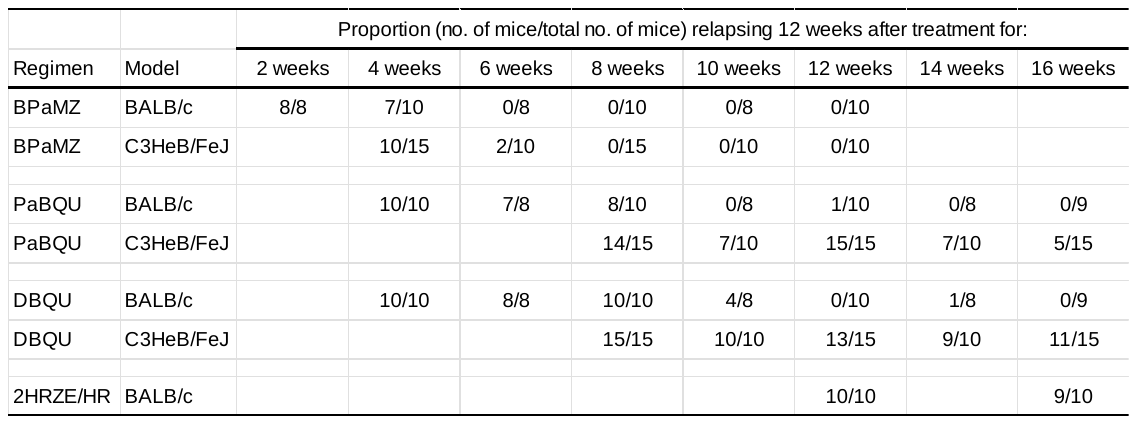

### **SECTION 1.** Sensitivity study of 5/7 versus 7/7 day dosing in the BALB/c model

**Purpose**.

This manuscript leveraged results from separately designed relapse studies. There was a difference in which DBQU and PaBQU were dosed 7 days a week in the BALB/c study, but only 5 of 7 days per week in the C3HeB/FeJ study. To evaluate the degree to which this may have biased our results, we conducted a 1-month sensitivity study comparing 5/7 versus 7/7 day doing in the BABL/c model.

**Methods**.

**Murine Infection Models.** All procedures and protocols for infecting mice with Mtb and subsequent drug treatments were approved by the Colorado State University Institutional Animal Care and Use Committee (IACUC) (Approved protocol number: KP 5172). Mice were housed in a certified animal bio-safety level III (ABSL-3) facility. Water and mouse chow were provided ad libitum. Mice were block randomized after aerosol infection and distributed into the different treatment arms. Infected mice were observed daily and weighed up to two times per week, due to the increased incidence of morbidity and mortality associated with clinical TB disease in the subacute BALB/c model. Any mice exhibiting clinical symptoms of illness were humanely euthanized.

**Drugs and experimental compounds.** Quabodepistat and spray-dried delamanid were provided by Otsuka. Bedaquiline Fumarate was obtained through the NIH HIV Reagent Program, Division of AIDS, NIAID, NIH: Bedaquiline Fumarate, ARP-12702, contributed by Janssen Pharmaceuticals. All other drugs were from commercial sources.

**Drug formulation.** Delamanid (D) and quabodepistat (Q) were formulated in 5% gum Arabic. Sutezolid (U) was formulated in 10% polysorbate 80, 40% polyethylene glycol 400 solution. Bedaquiline fumarate (B) was formulated in an acidified 20% 2-hydroxypropyl-β-cyclodextrin solution. Pretomanid (Pa) was formulated in a 10% 2-hydroxypropyl-β-cyclodextrin solution. Moxifloxacin hydrochloride (M) and pyrazinamide (Z) were formulated in sterile water.

**BALB/c subacute TB high dose aerosol model.** Female 6- to 8-week-old BALB/c mice were from Jackson Laboratories (Bar Harbor, ME). Mice were aerosol-infected using an inhalation exposure system (Glas-Col, Terre Haute, IN) with Mtb Erdman sub-cultured in Middlebrook 7H9 broth supplemented with 0.2% glycerol, 10% acid-albumin-dextrose-catalase (ADC) and 0.05% Tween 80 and adjusted to an optical density at 600 nm (OD600) of 1.0. This resulted in the average deposition of 4.16 log10 CFU in lungs of one-day after aerosol. Six mice were humanely killed 14 days post aerosol (D0) to determine CFU lung burdens at the start of treatment, resulting in an average 7.32 log10 CFU lung burden. The remaining mice received one of 5 different treatments, BPaMZ (5/7), PaBQU (5/7), PaBQU (7/7), DBQU (5/7), and DBQU (7/7), with the days dosed per week in parenthesis, respectively. With combinations of drugs at the following doses (in mg/kg body weight): B (25), Pa (50 [PaBQU] or 100 [BPaMZ]), D (6), Q (9), M (100), U (50), Z (150). All treatments were divided into 2 separate daily doses administered by oral gavage in 0.2 mL volume with > 1 h between doses. For BPaMZ, B was combined with Pa (1st dose), and M was combined with Z (2nd dose). For PaBQU and DBQU, either Pa or D was administered alone (1st dose), and B was combined with Q and U (2nd dose). BPaMZ was included as a reference benchmark control and was administered 5 of 7 days per week.

**Drug efficacy experiments.** Efficacy determinations were based on lung CFU counts from the left, caudal (inferior), and accessor (post-caval) lobes of the lung. The superior and middle lobes were collected for RNA. Tissues were aseptically harvested and frozen in 7mL Bertin Precellys tubes (CKmix50_7mL P000939-LYSK0A); tissues for RNA were flash frozen in liquid nitrogen. Samples for CFU were thawed, homogenized (Precellys, Bertin Instruments, Rockville, MD) and serially diluted in PBS with 10% [w:v] bovine serum albumin. Portions of the homogenates from animals on treatment were plated for CFU on 7H11-OADC agar (i.e., Middlebrook 7H11 agar plates supplemented 0.2% [v:v] glycerol, 10% [v:v] oleic acid-albumin-dextrose-catalase (OADC) supplement, cycloheximide (10 mg/L), carbenicillin (50 mg/L), amphotericin B (10 mg/L), polymyxin B (25 mg/L), trimethoprim (20 mg/L) and further supplemented with 0.4% [w:v] activated charcoal (7H11 charcoal agar) to help counteract drug carryover artifacts. Colonies were enumerated after at least 28 days of incubation at 37°C and plates were incubated for ≥ 6 weeks to ensure all viable colonies were detected. Mice were euthanized by CO_2_ inhalation followed by cervical dislocation, a method approved by the IACUC at Colorado State University.

**RNA extraction.** RNA was extracted from superior and middle lung lobes in CAMM-RPS, as described previously (1).

**RS ratio.** Details on the RS ratio pharmacodynamic measure were described previously (2).

**Results**.

**Figure S1.** Longitudinal RS ratio and CFU results in the BALB/c high-dose aerosol model with DBQU dosed 5/7 (light green) versus 7/7 (dark green) days per week. Dots represent values from individual mice. Lines connect mean values.

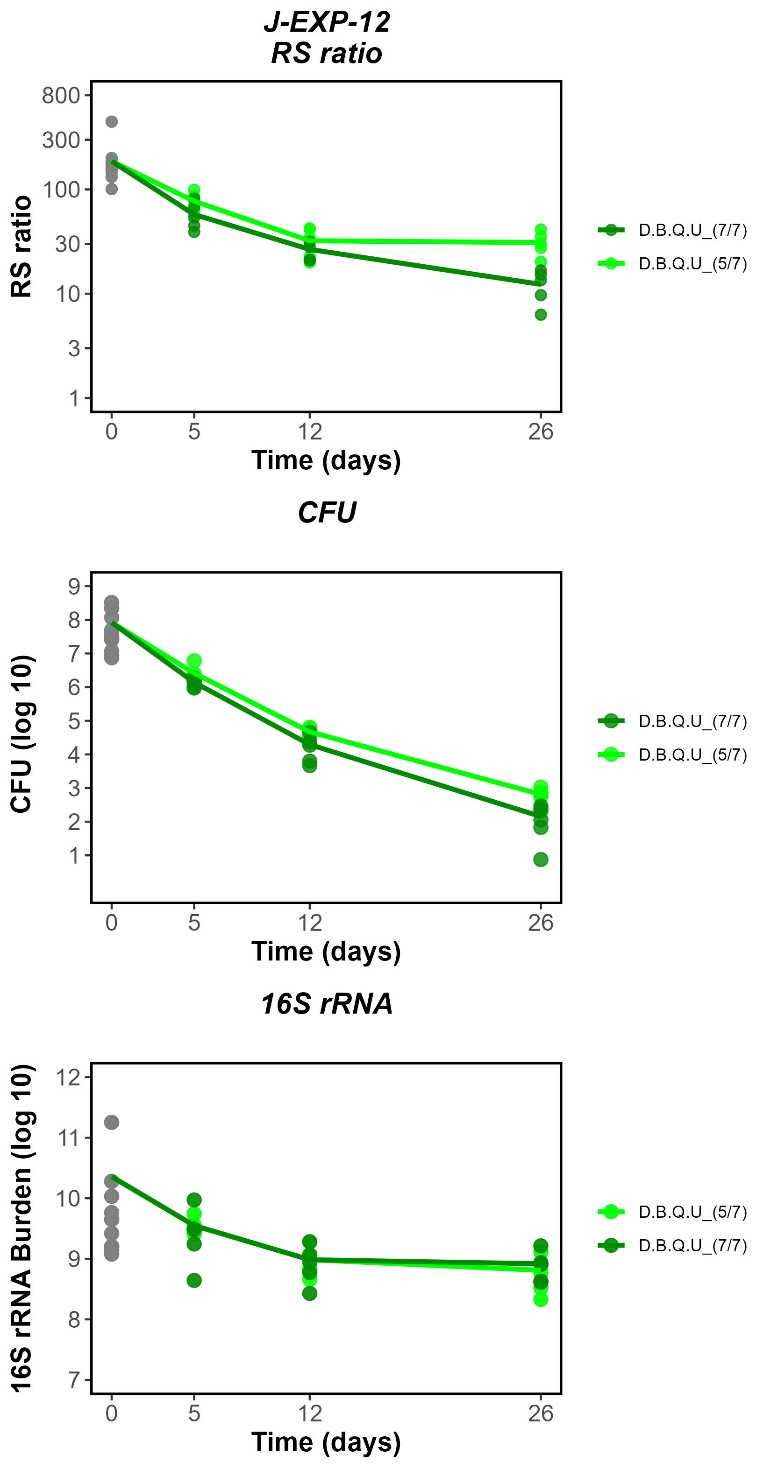

**Figure S2.** Longitudinal RS ratio and CFU results in the BALB/c high-dose aerosol model with PaBQU dosed 5/7 (light red) versus 7/7 (dark red) days per week. Dots represent values from individual mice. Lines connect mean values.

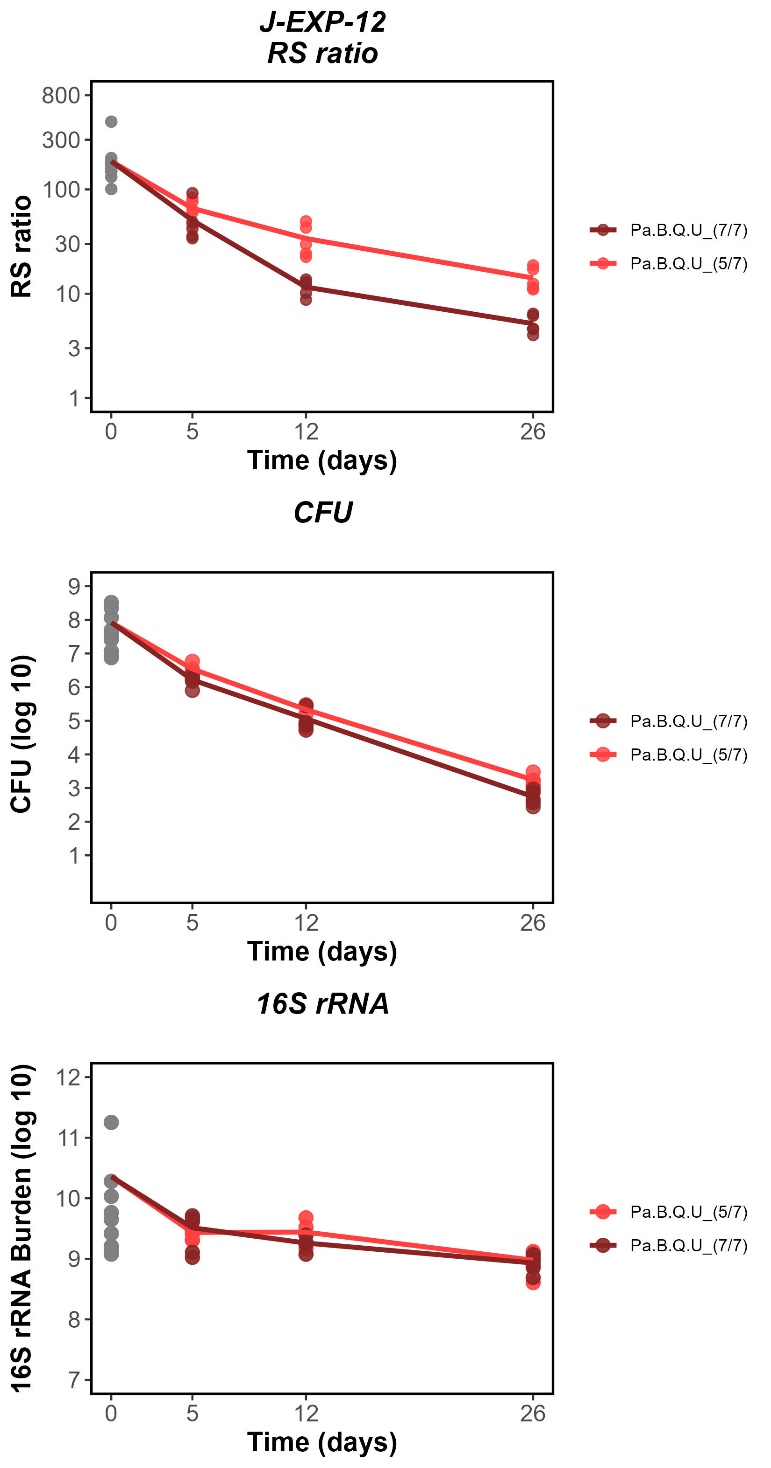

**Conclusion of 5/7 versus 7/7 day dosing sensitivity study**. The bactericidal activity of DBQU or PaBQU did not differ between mice dosed 5/7 versus 7/7 days per week. This indicates that the difference in dosing schedule does not explain the lesional liability in bactericidal activity observed between BALB/c and C3HeB/FeJ mice. The RS ratio activity of DBQU was marginally greater on day 26 among mice dosed 7/7 versus 5/7 days/week and the RS ratio activity of PaBQU was greater on days 12 and 26 among mice dosed 7/7 days/week. Because the scale of the difference between 5/7 and 7/7 dosing in BABL/c mice is much smaller than the RS ratio difference between BALB/c and C3HeB/FeJ, we conclude that this does not affect our central findings regarding lesional liability.

**Acknowledgements.**

The following reagent was obtained through the NIH HIV Reagent Program, Division of AIDS, NIAID, NIH: Bedaquiline Fumarate, ARP-12702, contributed by Janssen Pharmaceuticals

**References.**

1.	Wynn EA, Dide-Agossou C, Reichlen M, Rossmassler K, Al Mubarak R, Reid JJ, Tabor ST, Born SEM, Ransom MR, Davidson RM, Walton KN, Benoit JB, Hoppers A, Loy DE, Bauman AA, Massoudi LM, Dolganov G, Strong M, Nahid P, Voskuil MI, Robertson GT, Moore CM, Walter ND. 2023. Transcriptional adaptation of *Mycobacterium tuberculosis* that survives prolonged multi-drug treatment in mice. mBio 14:e0236323.

2.	Walter ND, Born SEM, Robertson GT, Reichlen M, Dide-Agossou C, Ektnitphong VA, Rossmassler K, Ramey ME, Bauman AA, Ozols V, Bearrows SC, Schoolnik G, Dolganov G, Garcia B, Musisi E, Worodria W, Huang L, Davis JL, Nguyen NV, Nguyen HV, Nguyen ATV, Phan H, Wilusz C, Podell BK, Sanoussi ND, de Jong BC, Merle CS, Affolabi D, McIlleron H, Garcia-Cremades M, Maidji E, Eshun-Wilson F, Aguilar-Rodriguez B, Karthikeyan D, Mdluli K, Bansbach C, Lenaerts AJ, Savic RM, Nahid P, Vásquez JJ, Voskuil MI. 2021. *Mycobacterium tuberculosis* precursor rRNA as a measure of treatment-shortening activity of drugs and regimens. Nat Commun 12:2899.
